# Descending pathways from the lateral accessory lobe and posterior slope in the brain of the silkmoth *Bombyx mori*

**DOI:** 10.1101/247619

**Authors:** Shigehiro Namiki, Ryohei Kanzaki

**Affiliations:** Research Center for Advanced Science and Technology, The University of Tokyo, 4-6-1 Komaba, Meguro, Tokyo 153-8904, Japan

**Keywords:** Intracellular recording, lateral accessory lobe, posterior slope, pheromone orientation, thoracic ganglion

## Abstract

A population of descending neurons connect the brain and thoracic motor cener, playing a critical role in controlling behavior. We examined the anatomical organization of descending neurons (DNs) in the brain of the silkmoth *Bombyx mori*. Moth pheromone orientation is a good model to investigate the neuronal mechanisms of olfactory behavior. Based on mass staining and single-cell staining, we evaluated the anatomical organization of neurite distribution by DNs in the brain. Dense innervation was observed in the posterior–ventral part of the brain, called the posterior slope (PS). We examined the morphology of DNs innervating the lateral accessory lobe (LAL), which is assumed to be important for moth olfactory behavior. We observed that the LAL DNs also innervate the PS, suggesting the integration of signals from the LAL and PS. We also identified a set of DNs innervating the PS, but not the LAL. These DNs were sensitive to sex pheromones, suggesting a role of the PS in motor control for pheromone orientation. The organization of descending pathways for pheromone orientation is discussed.

## 1. Introduction

Male moths orient toward conspecific females based on their sex pheromones. Sex pheromones have been shown to reliably elicit stereotyped behavior in moths and, hence, can be used as a good model for investigating the general mechanisms underlying olfactory navigation ^1^. A specific group of descending neurons (DNs), which connect the brain and ventral nervous system, have been identified as an important element for pheromone orientation in the silkmoth ^2^. These DNs have neurites innervating a specific brain region called the lateral accessory lobe (LAL), which is a key player in insect brains connecting the central complex with other parts of the protocerebrum ^3,4^. Only a fraction of DNs innervate the LAL and the individual morphology is largely unknown for DNs innervating other regions of the brain. Insects have 300– 500 DNs from each hemisphere ^5–8^. Recent studies have reported that the number of DNs from the LAL is smaller than from other brain regions in *Drosophila* ^9^.

The posterior ventral part of the brain can be densely labeled using backfill labeling from the neck connective in insects ^6,7,10^. A systematic analysis of the individual DN morphology revealed that the largest number of DNs originate from the area known as the posterior slope (PS) in *Drosophila melanogaster* ^9^. The PS is densely labeled using backfill staining, suggesting that the PS is the major origin of DNs in the silkmoth *Bombyx mori*, as in other species ^11^. The neuroanatomy of DN innervation in the PS is, however, largely unknown in the silkmoth and the functional role on pheromone orientation remains unclear. Here, we investigate the morphology of individual DNs in the silkmoth by using single-cell and backfill labeling techniques. We describe the gross anatomical organization of the descending pathway and particularly focus on the innervation of the PS.

In a previous study, we classified DNs into three groups based on the cell body location (Fig. 1) ^2^. Group-I and -II DNs, whose cell bodies are located on the anterior surface (Fig. 1) and group-III DNs, whose cell bodies are located on the posterior surface have been identified thus far in *B. mori* ^2,12^. The pheromone-evoked flip-flop firing activity, which correlates with the walking direction during pheromone orientation ^13^, has been observed in the neurons innervating the LAL ^11,14^. In the present study, we found that these LAL DNs also have neurite innervation in the PS as well as in the LAL. In addition, we identified several types of group-III DNs innervating the PS. The organization of descending pathway is discussed based on the neuroanatomy.

**FIGURE 1.**
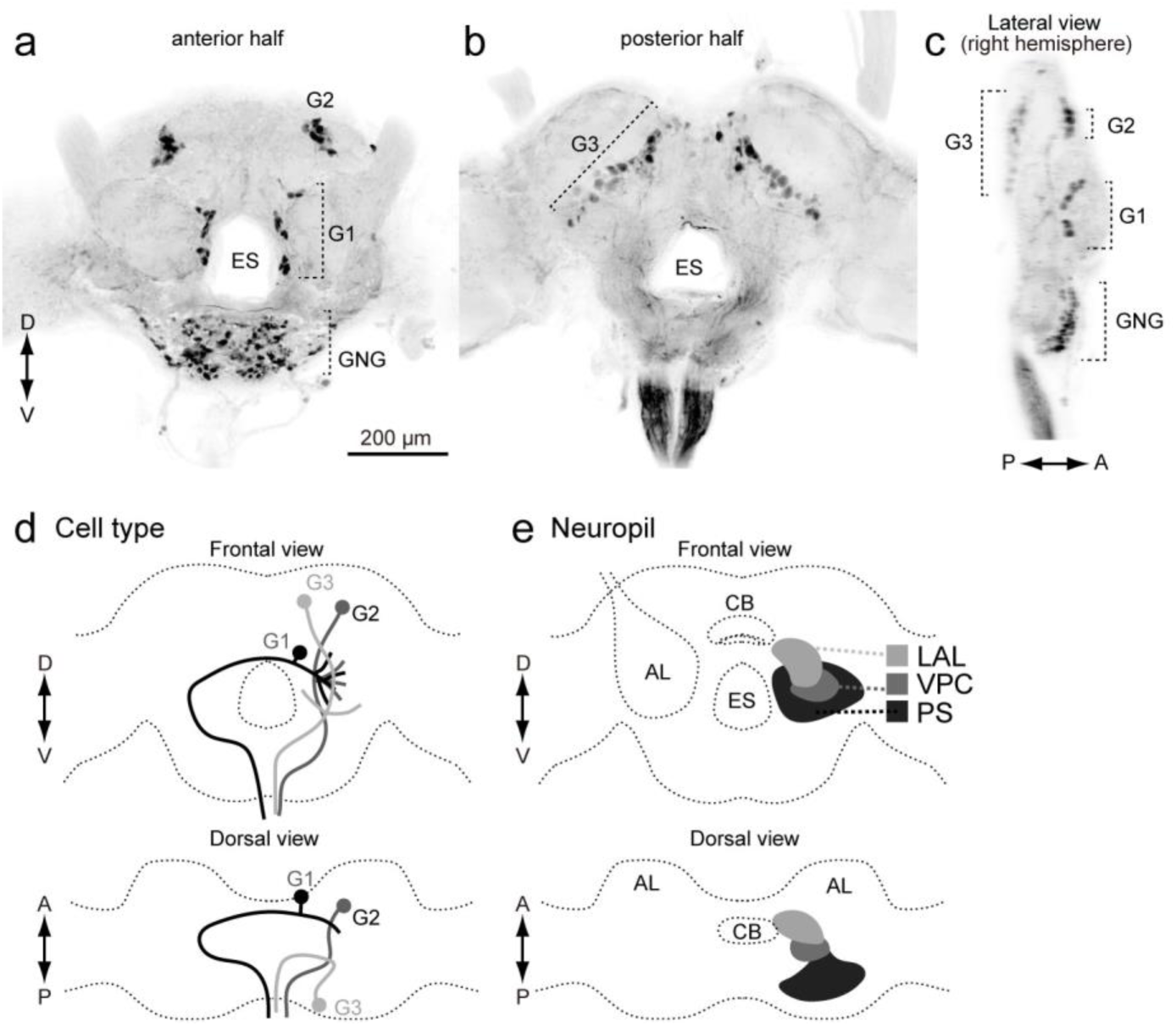
Basic anatomy of descending neurons and major innervation area in the silkmoth brain. (**a,b**) Maximum intensity projection of anterior (a) and posterior part of brain sample stained by backfilling technique (b). (**c**) Lateral view of the brain sample shown in (a,b). (**d**) Classification of DNs based on the cell body location. (**e**) Schematic of neuropils in the silkmoth brain. Frontal and dorsal views are shown. AL, antennal lobe; CB, central body; ES, esophagus; G1, group-I DNs; G2, group-II DNs; G3, group-III DNs; GNG, gnathal ganglion; LAL, lateral accessory lobe; PS, Posterior slope; VPC, ventral protocerebrum.

## 2. Results

### 2.1. Organization of descending neurons in the brain

To characterize the anatomical organization of DNs, we performed backfill labeling from the neck connective using florescent dye (Fig. 2). Figure 2b shows images of staining at three depths, corresponding to the LAL, ventral protocerebrum (VPC), and PS of the left hemisphere. The posterior part of the protocerebrum, including the PS, was more densely labeled than the anterior anterior part, including the LAL and VPC (Fig. 2c; Supplementary Fig. 1). Furthermore, the posterior lateral protocerebrum (PLP) and ventral lateral protocerebrum (VLP), which mainly receives input from the lobula complex and inferior bridge (IB) located beside the protocerebal bridge, were densely labeled (Supplementary Fig. 2). The innervation to the PLP was further extended to the lobula (Supplementary Fig. 2b). We then analyzed the innervation within the LAL, a circuit known to be important for pheromone orientation ^11^. The LAL is classified into two subdivisions roughly delineated by the lateral accessory lobe commissure (LALC). Backfilling labeled more innervation in the lower division than in the upper division (Fig. 2d).

**FIGURE 2.**
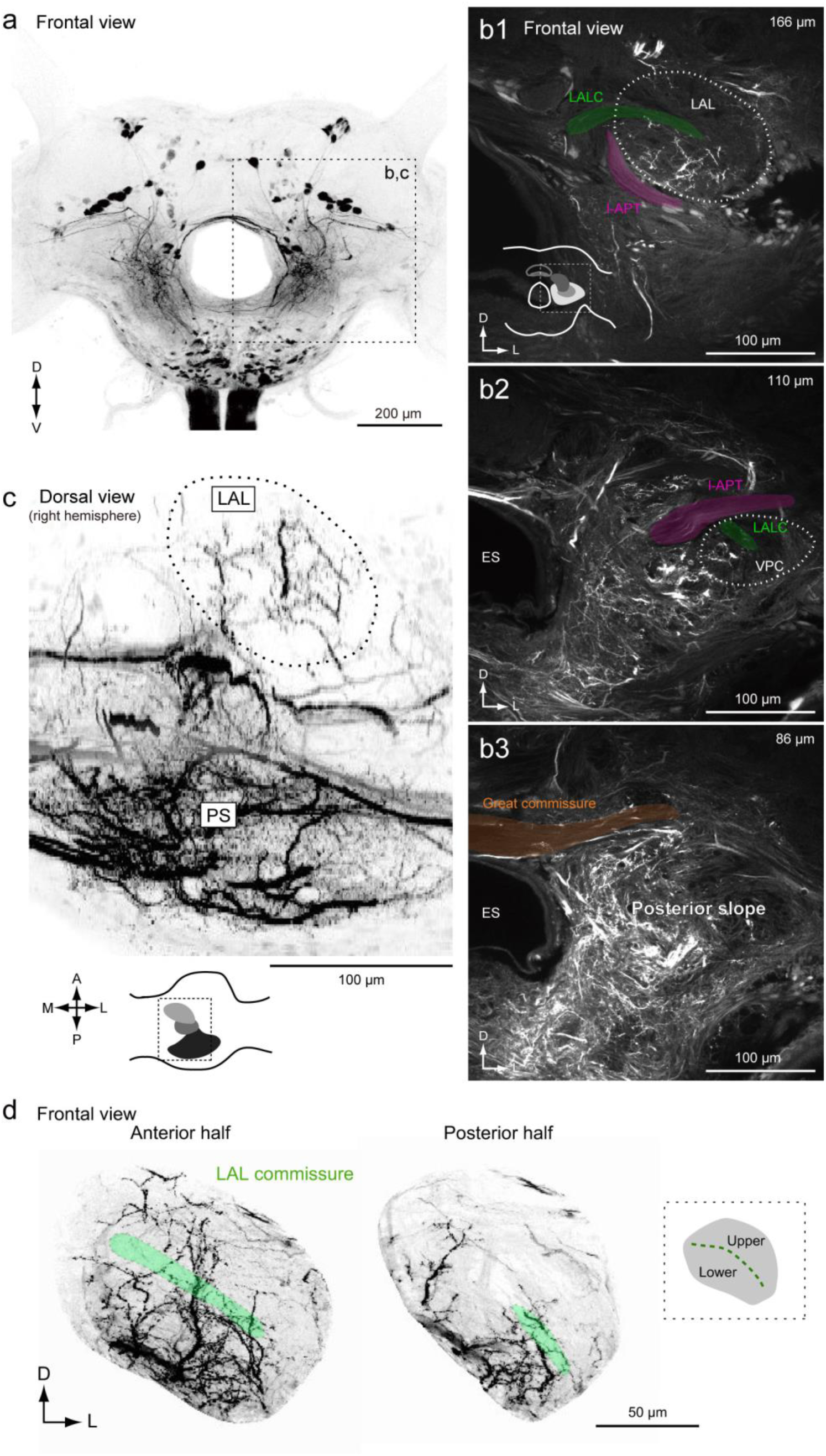
Anatomical organization of descending and ascending neuron innervation in the brain. (**b**) Mass-staining results by backfilling from the neck connective on both sides. Frontal and lateral views are shown. The positions of DN groups are shown (*right*). G1, group-I DNs; G2, group-II DNs; G3, group-III DNs; GNG, DNs of gnathal ganglion. (**c**) Confocal stacks of mass-staining result in the protocerebrum. The depth from the posterior brain surface are shown in *top-right*. Inset shows the schematic of the imaging area (c1). The shape of the LAL and VPC are shown with broken line. The location of major bundles are shown with color: the lateral accessory lobe commissure (LALC, green), lateral antennal-lobe tract (l-ALT, magenta) and great commissure (GC, orange). (**d**) Dorsal view of innervation of descending and ascending neurons. Inset shows the imaging area. The shape of LAL is shown with broken line. The innervation in the PS is more dense than in the LAL. (**e**) Neurite innervation in the LAL. The maximum intensity projection for the anterior and posterior half of the LAL are shown. The position of the LALC is indicated by green. Inset shows subdivisions within the LAL. The upper and lower divisions are roughly delineated by the LALC. The innervation is observed in both upper and lower divisions in the anterior part, whereas largely confined to the lower division in the posterior part of LAL.

To compare the DN innervation of the ipsilateral and contralateral sides, we examined backfilling from one side of the neck connective (Fig. 3). This resulted in bilateral labeling of the PS, indicating both ipsilaterally and contralaterally descending fibers. Ipsilateral innervation was broader than contralateral innervation (Fig. 3b, c). The contralateral innervation was biased toward the posterior side and innervation of the anterior side was rare (Fig. 3d). Biased innervation within the LAL (Fig. 2d) was also confirmed when we filled one side of the neck connective (Supplementary Fig. 3).

**FIGURE 3.**
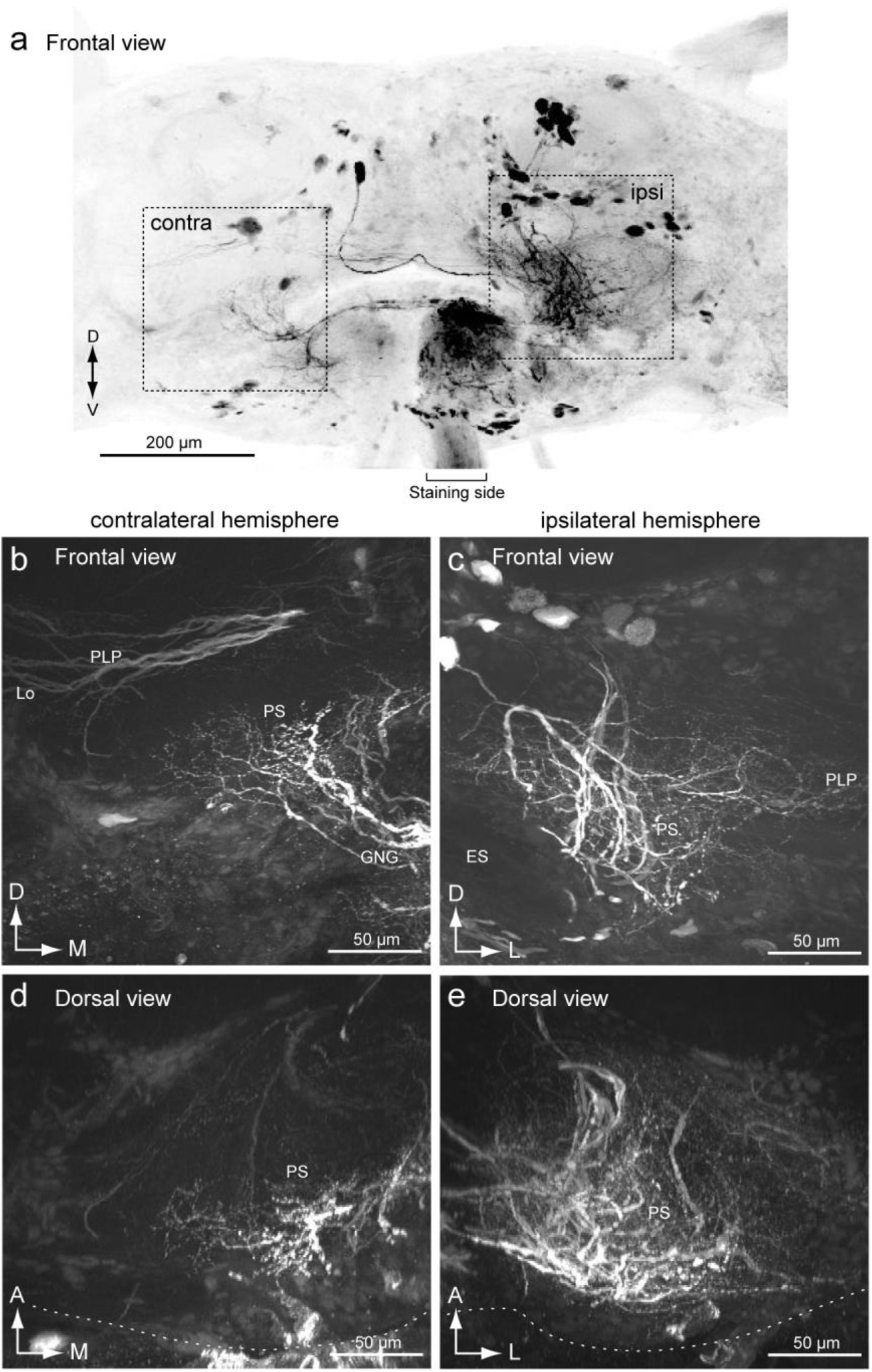
Mass-staining result by backfilling from one side of the neck connective. (**a**) Maximum intensity projection of mass-staining result. Frontal view is shown. The side of filling is noted. (**b,c**) Frontal view of the innervation in the posterior slope (PS) of contralateral (b) and ipsilateral side (c). Wide field innervation is observed in the ipsilateral hemisphere. (**d,e**) Dorsal view of the innervation in the PS of contralateral (d) and ipsilateral side (e). The innervation is extended toward more anterior part in the ipsilateral hemisphere. ES, esophagus; GNG, gnathal ganglia; PLP, posterior lateral protocerebrum.

Furthermore, to characterize the anatomical organization at a single cell resolution, we analyzed the morphology of 57 DNs (Supplementary Table. 1). Among them, 38 and 19 were ipsilaterally and contralaterally descending, respectively, and 22 had branches crossing the midline. Most DNs had varicose processes in the gnathal ganglion (GNG) (98%, *n* = 57) and 22 DNs had varicose processes in the PS (39%, *n* = 57). The PS was the region that contained the largest number of DNs in the brain (86%, *n* = 57). Almost all DNs supply varicose processes in the GNG in the silkmoth (Supplementary Table 1), suggesting that most DNs provide an output signal to both the ventral nervous system and gnathal ganglion. This anatomical feature is observed in flies where 78% of DN types have output terminals in the GNG ^9,15^. Some group-I and II DNs may have monosynaptic contact on the neck motor neurons ^12^. We describe their morphology as follows.

### 2.2. Group-I descending neurons

We analyzed a total of 11 group-I DNs (Supplementary Table. 1). We observed smooth processes in the PS for all DNs. Figure 4 shows the morphology of all three types of DNs identified in a previous study ^2,12^. Most of them were contralaterally descending (91%, n = 11). Most of the group-I DNs have smooth processes in the LAL and PS of the ipsilateral hemisphere and varicose processes in the PS and GNG of the contralateral hemisphere. Group-IA has smooth processes in the small portion of the medial PS of the ipsilateral hemisphere and varicose processes in part of the medial PS of the contralateral hemisphere (Fig. 3a1; Supplementary Fig. 4). Group-IB has smooth processes in the medial PS of the ipsilateral hemisphere, over a wider area than do Group-IA (Fig. 3a2; Supplementary Fig. 5). Group-IB has varicose processes in the medial PS of the contralateral hemisphere, different from Group-IA. In contrast, the smooth processes of GIC occupy the entire volume of the PS of the ipsilateral hemisphere (both medial and lateral PS) and while varicose processes occupy the PS of the contralateral hemisphere, which is similar to those of Group-IB (Fig. 3a3; Supplementary Fig. 6). A schematic of the group-I DN’s neurite innervation is shown in Figure 4c. The smooth innervation shows a differential size of arborization and the varicose processes exhibit a bias to innervating the medial PS.

**FIGURE 4.**
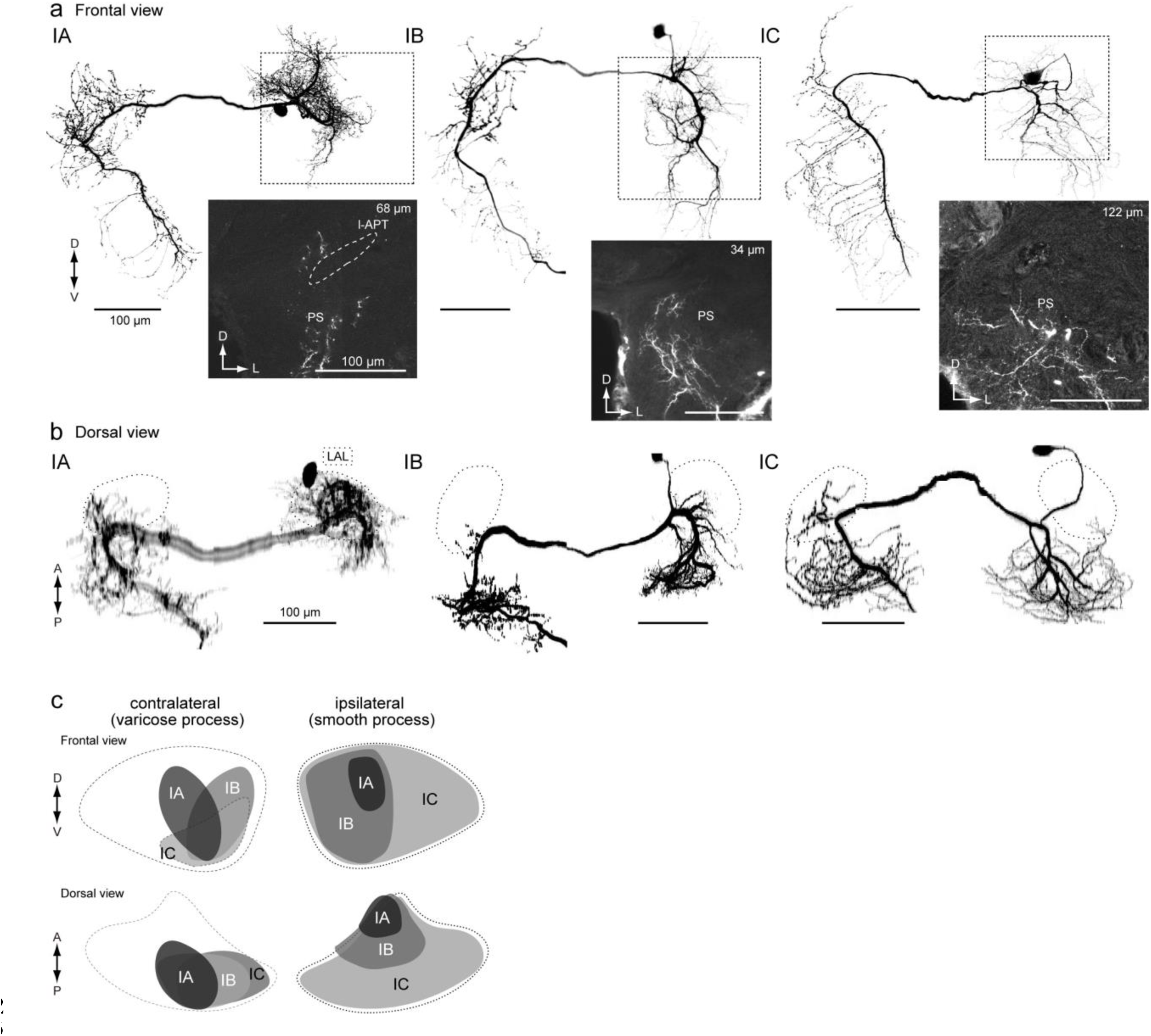
Neurite innervation to the posterior slope PS by group-I and group-II descending neurons. (**a**) Morphology of group-I DNs. Maximum intensity projection of whole innervation in the brain(*top*) and confocal stack of the neurite innervation in the posterior slope (PS) are shown (*bottom*). Frontal view is shown. GIA, GIB and GIC innervate a part of medial PS, medial PS and entire PS, respectively. The original data are taken from 12 for GIB. (b) Dorsal view of DN morphology. Shape of the lateral accessory lobe (LAL) is shown with broken line. There are dense innervation outside the LAL. (**c**) Schematic for innervation of group-I DNs in the PS. Shape of the PS is shown with broken line. The innervation of varicose process is biased toward the medial side (*left*). The innervation of smooth process differs across DN types (*right*). l-APT, lateral antennal-lobe tract.

In addition to the types described so far, we identified novel types of group-I DNs whose cell bodies are located on the anterior brain surface (Figs. 5 and 6). These DNs may correspond to group-I’ DNs, whose cell body position is located on the anterior brain surface and close to group-I cell group ^11^. Figure 5 shows the morphology of unilateral DN. The DN has smooth processes in the LAL, VPC, and PS and varicose processes in the PS and GNG of the ipsilateral hemisphere. Thin neurites extends to the PLP and inner lobula (Fig. 5a). The innervation within the LAL was biased toward the lower division (Fig. 5c), similar to the other types of group-I DNs. Unlike other group-I DNs, this DN descends to the ipsilateral neck connective.

**FIGURE 5.**
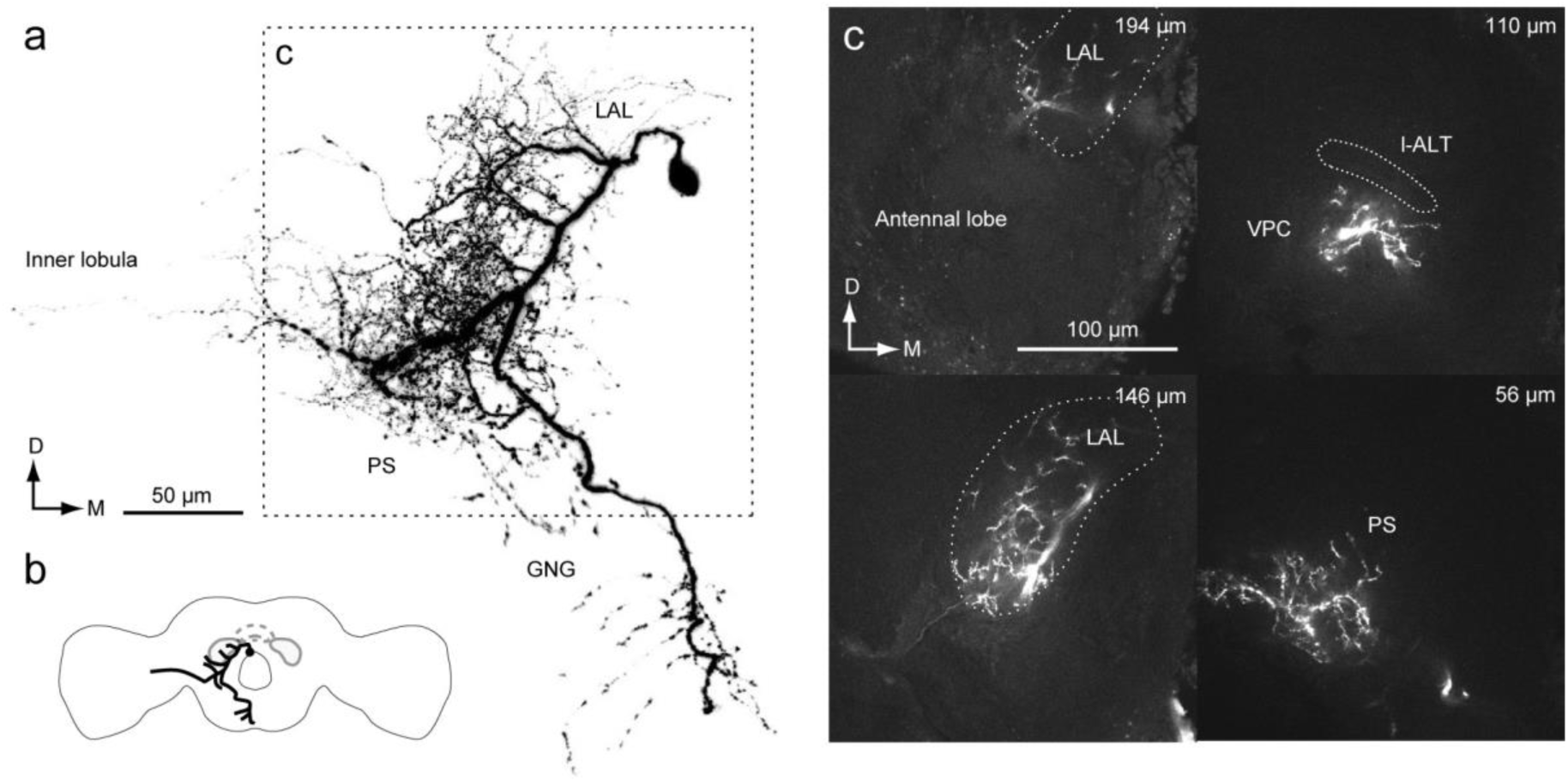
Newly identified group-I DN innervating the LAL. (**a**) Whole morphology of the DN. The DN has smooth processes in the ipsilateral lateral accessory lobe (LAL), ventral protocerebrum (VPC) and posterior slope (PS) and varicose processes in the ventral PS and GNG. The axon runs thought the medial route in the GNG and descends ipsilateral neck connective. (**b**) Schematic of DN morphology. (**c**) Confocal stacks of neurite innervation. The depth from the posterior brain surface is shown in *top-right*. The innervation is biased toward the lateral side within the LAL. The innervation in the PS is biased toward the lateral side.

Figure 6 shows another example of group-I DN. The DN has smooth processes ranging a wide area in the protocerebrum, PS, and IB, and varicose processes in the GNG. Some neurites in the IB crosses the midline. The lateral branch reaches the PLP. Unlike the other group-I DNs, this DN does not innervate the LAL.

**FIGURE 6.**
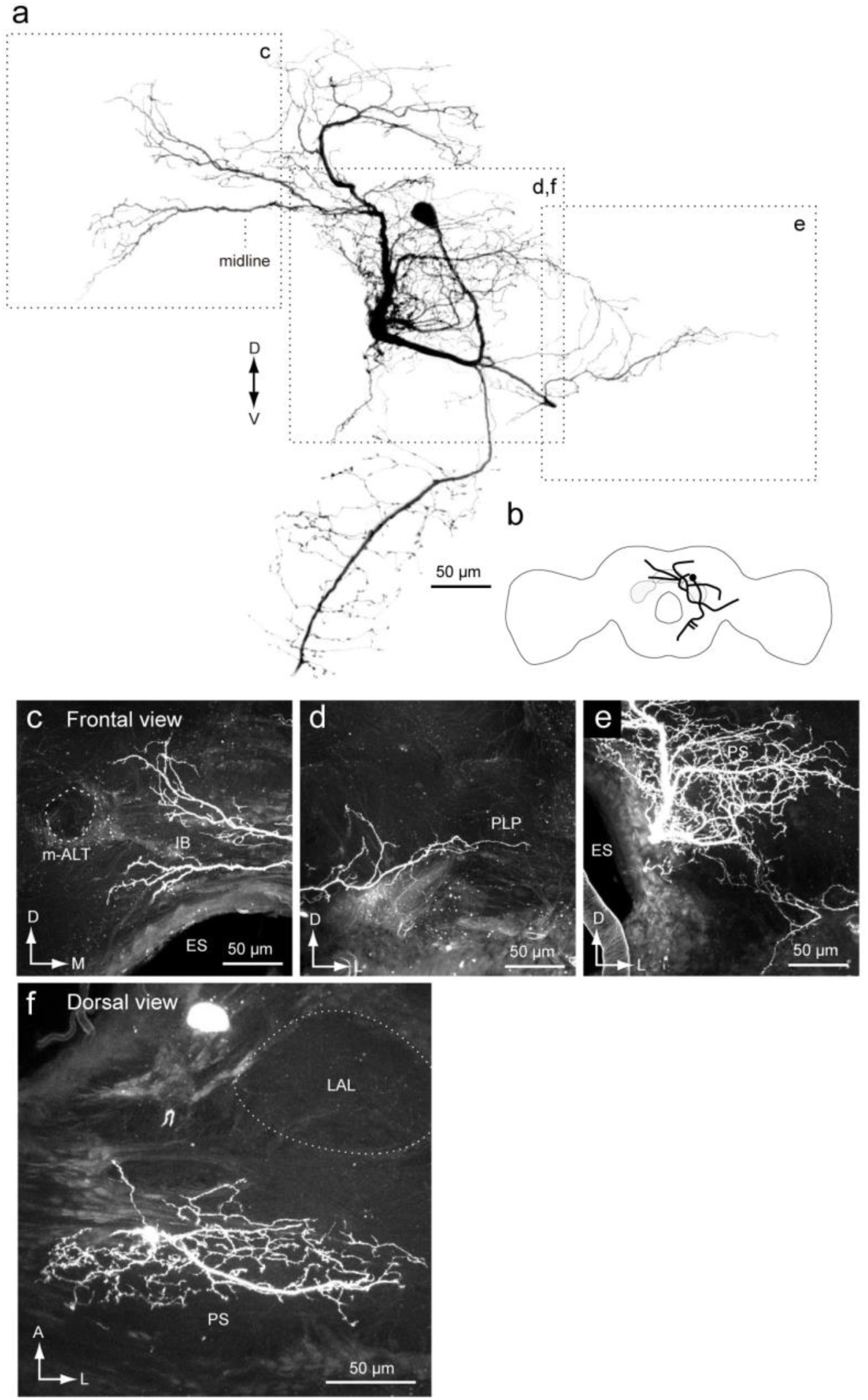
Newly identified group-I DN. (**a**) Whole morphology of the DN. The (**b**) Schematic of theneuronal morphology. DN has smooth processes in the inferior bridge (IB) (**c**), posterior lateral protocerebrum (PLP) (**e**) and the posterior slope (PS) (**e**). Unlike other group-I DNs, the DN does not have innervation to the lateral accessory lobe (LAL) (**f**). ES, esophagus; m-ALT, medial antennal-lobe tract.

### 2.3. Group-II descending neurons

We analyzed 29 group-II DNs, which are classified into a total of six cell types (group-IIA–F, Supplementary Table 1). All types are ipsilaterally descending and had smooth processes in the PS of the ipsilateral hemisphere and varicose processes in the GNG of the contralateral hemisphere. Figure 7 shows the morphology of group-II DNs. All DN types have innervations in the VPC and PS.

**FIGURE 7.**
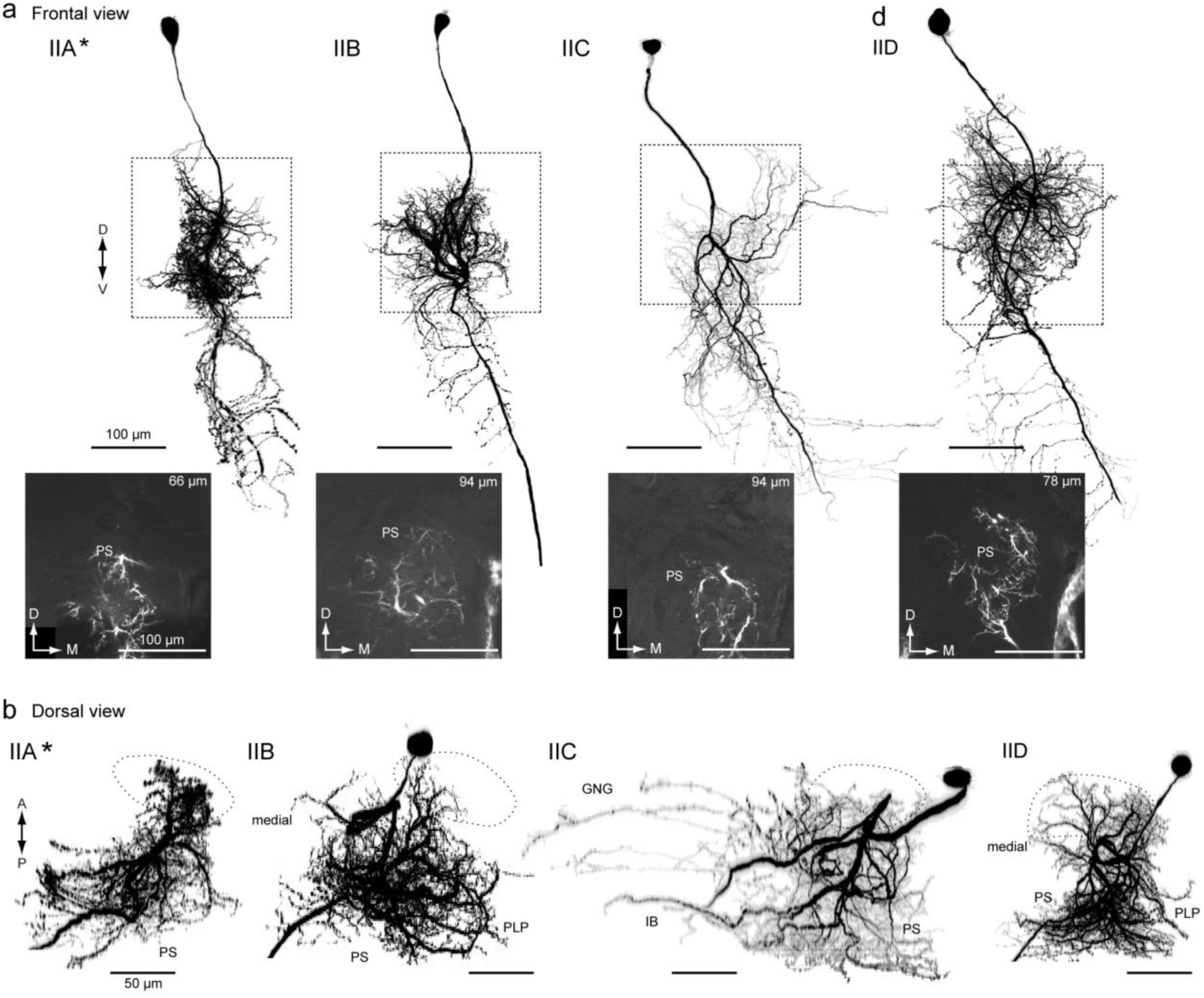
Group-II DNS iinnervates the PS. (**a**) Morphology of group-II DNs. Maximum intensityprojection of whole innervation in the brain (*top*) and confocal stack of the neurite innervation in the posterior slope (PS) are shown (*bottom*). Frontal view is shown. Flipped image is shown for the purpose of comparison (asterisk). All DN types have innervation in the medial PS. (**b**) Drosal view of DN innervation in the brain. The shape of the LAL is shown with broken line. The innervation area are similar among group-II DNs, in comparison with group-I DNs (Fig. 4). GNG, gnathal ganglion; IB, inferior bridge; PLP, posterior lateral protocerebrum; PS, posterior slope.

Group-IIA innervates the LAL, VPC, and medial part of the PS, which is located medial to the LALC. Moreover, the DN has a number of branches entering the unstructured region medial to the LAL. These branches run dorsal and ventral to the l-ALT (Supplementary Fig. 7a, asterisks). In the deep PS, the innervation is biased toward the dorsal side and lacking in the ventral side. The ventral branch is bifurcated and enters the GNG.

Group-IIB innervates the LAL, VPC, and medial part of the PS (Supplementary Fig. 7b). As with group-IIA, the DN had branches in the ventral end of the superior medial protocerebrum (SMP), and the innervation was biased to the dorsal aspect of the posterior side. The area innervated in the PS is wider than for the other types. The DN has smooth branches located posterior to the LAL and dorsal to the lateral antennal-lobe tract (l-ACT). The axon runs through the medial route of the GNG with side branches.

Group-IIC innervates the VPC and medial side of the PS. The DN has additional processes in the posterior medial part of the protocerebrum, called the IB (Fig. 7, Supplementary Fig. 7c), which is posterior of the midline region behind the central body upper division and below the protocerebral bridge ^16,17,18^. The medial branch into the IB often crosses the midline. The axon runs through the medial route in the GNG and side branches extend laterally; there are fewer side branches than in the other types.

Group-IID provided innervates to the LAL, VPC, and medial side of PS (Supplementary Fig. 7d). The innervation in the LAL is mostly localized to its lower division and the anterior portion of the upper division. Some processes reach the ventral end of the SMP. Some branches invaded the periesophageal area located medial to the LAL; the axons ran through the medial route in the GNG with many side branches toward the lateral side.

In addition, we identified two novel types of group-II DNs (Fig. 8). One shows similar morphology to the Group-IIC DN (Fig. 8a). The DN innervates the PS, but not the LAL or VPC. Some of the innervation supplies to the PS display a varicose appearance (Fig. 8c), which is likely to be postsynaptic ^19^. The DN is referred to as group-IIE DN. Another example, group-IIF DN shows similar morphology to group-IID (Fig. 8d). GIIF innervation of the PS is mostly smooth in appearance but also some shows varicose appearance (Fig. 8f).

**FIGURE 8.**
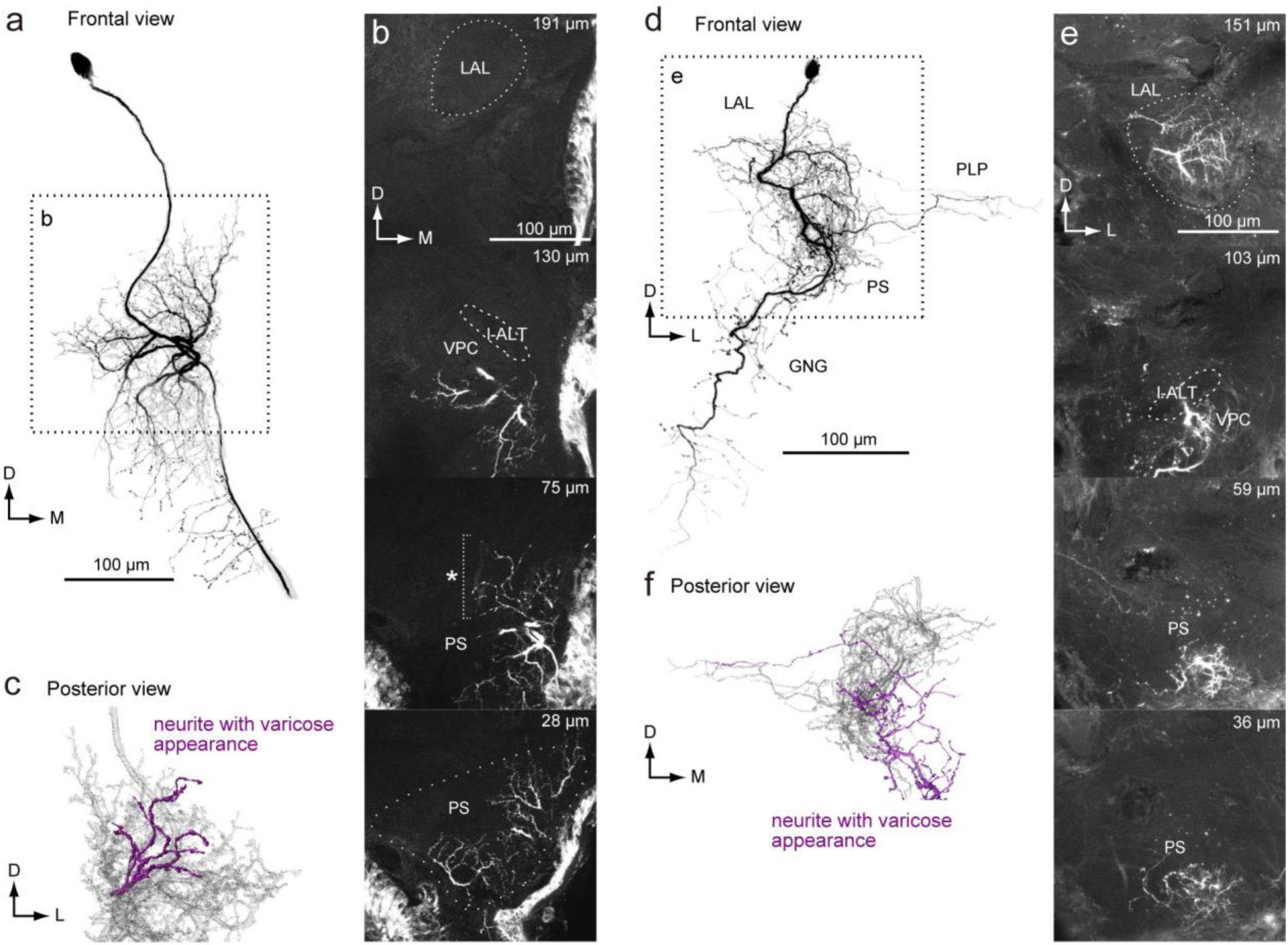
Newly identified group-II DNs. (**a**) Maximum intensity projection of the DN innervation inthe brain. The morphology is similar to group-IID DN. Unlike to other group-II DNs, the DN has innervation with varicose appearance in the PS. Also, the DN has innervation to the PLP. (**b**) Confocal stacks for the DNs in (a). The DN has innervation in the almost entire part of the LAL, ventral protocerebrum (VPC) and medial part of the inferior PS. The depth from the posterior surface are shown in *top-right*. (**c**) Posterior view of three-dimensional reconstruction of the DN in the PS. The neurite with varicose appearance are shown with magenta. (**d**) Maximum intensity projection of the DN innervation in the brain. The morphology is similar to group-IIC DN. Unlike to other group-II DNs, the DN has innervation with varicose appearance in the PS. (**e**) Confocal stacks for the DNs in (a). The DN has innervation in the almost entire part of the LAL, ventral protocerebrum (VPC) and inferior PS. The depth from the posterior surface are shown in *top-right*. (**f**) Posterior view of three-dimensional reconstruction of the DN in the PS. The neurite with varicose appearance are shown with magenta.

Overall, all group-II DN types have smooth processes in the medial side of the PS. All three types of GI-DNs have varicose processes in the medial PS of the contralateral hemisphere (Fig. 4c), overlapping with the smooth processes of group-II DNs, suggesting information flow from group-I to group-II DNs.

### 2.4. Group-III descending neurons

We identified a total of 17 group-III DNs (Supplementary Table. 1). Areas highly innervated by DNs included the PLP (10 DNs), PS (9), lobula (5), SMP (5), IB (4), and inferior clamp (ICL) (4) (Supplementary Figs. 8 and 9). Ten descended contralaterally, while seven descended ipsilaterally. As in group-I and group-II DNs, most group-III DNs had varicose processes in the GNG (94%, *n* = 17) and nine out of 17 DNs had varicose processes in the PS (53%).

Among these, four DNs innervate the PS but not the LAL (Figs. 9 and 10; Supplementary Figs. 10 and 11). Figure 9 shows an example of a DN innervating the PS, but not the LAL. The DN has smooth processes in the PS, PLP, and IB, and varicose processes in the GNG. The axon runs through the lateral route of the GNG. The DN exhibited an excitatory response to bombykol, the major sex pheromone, but not to bombykal, a behavioral antagonist of *B. mori* ^20^. Supplementary Figure 10 shows a neuron with the same morphology as Figure 9, which also responded to bombykol. Figure 10 shows another DN innervating the PS and GNG. The DN has smooth processes in the PS. The processes in the GNG mostly show a smooth appearance, but some neurites have a varicose appearance. The DN dose not innervate pheromone processing circuits, including the LAL ^21^, suggesting that the PS may also process sex pheromone information. Supplementary Figure 11 shows a neuron with the same morphology.

**FIGURE 9.**
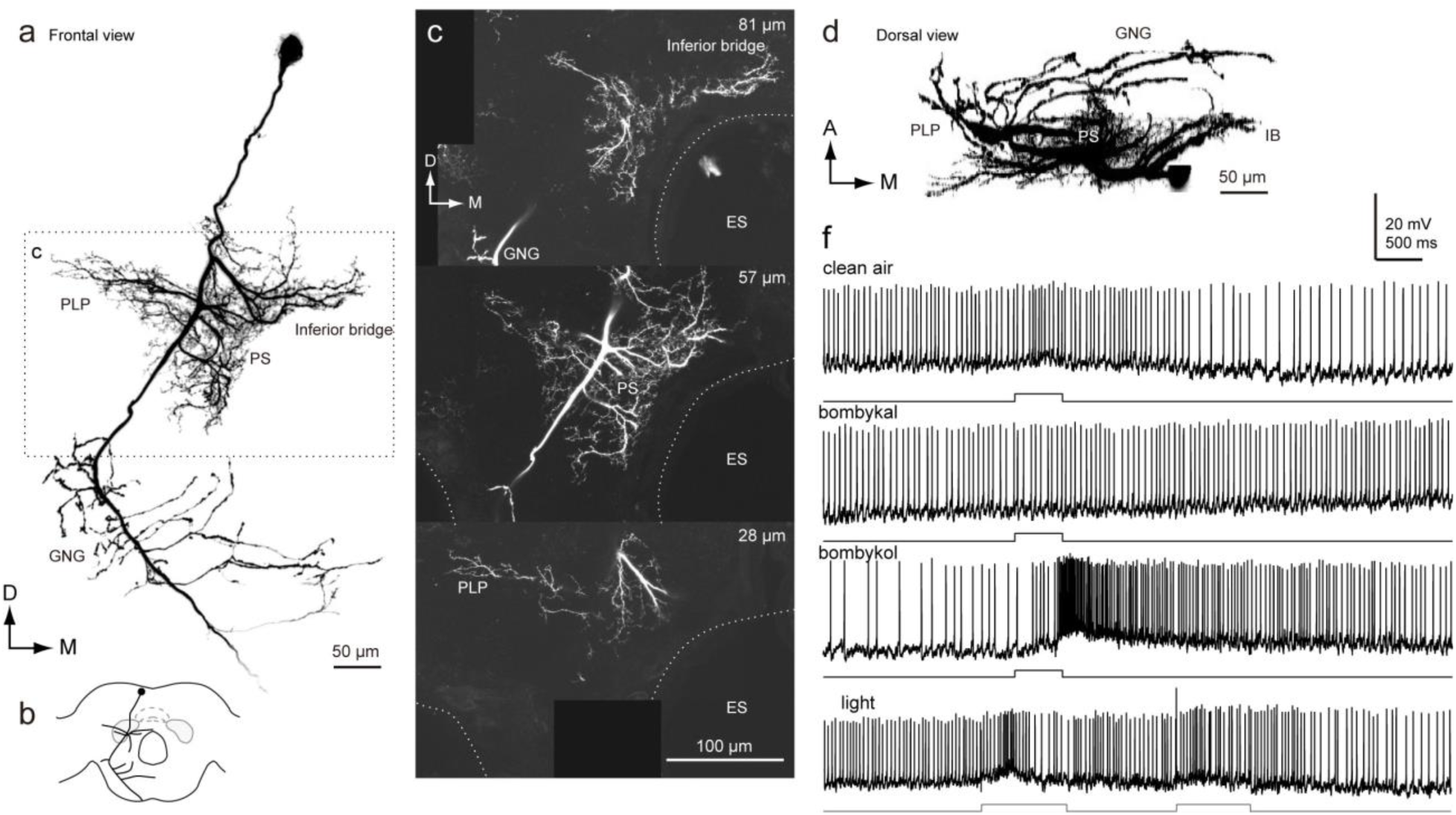
Morphology of a descending neuron innervating the PS. (**a**) Maximum intensity projection of the DN innervation in the brain. The neuron has smooth processes in inferior bridge (IB), posterior lateral protocerebrum (PLP) and posterior slope (PS), and varicose processes in the gnathal ganglion (GNG). (**b**) Schematic of neurite innervation for the neuron shown in (a). (**c**) Confocal stacks for the DNs shown in (a). The depth from the posterior brain surface is shown in *top-right*. (**d**) Dorsal view of the DN morphology. (**e**) Response property of the neuron. The neuron shows excitatory response to the exposure of bombykol, the sex pheromone. ES, esophagus.

**FIGURE 10.**
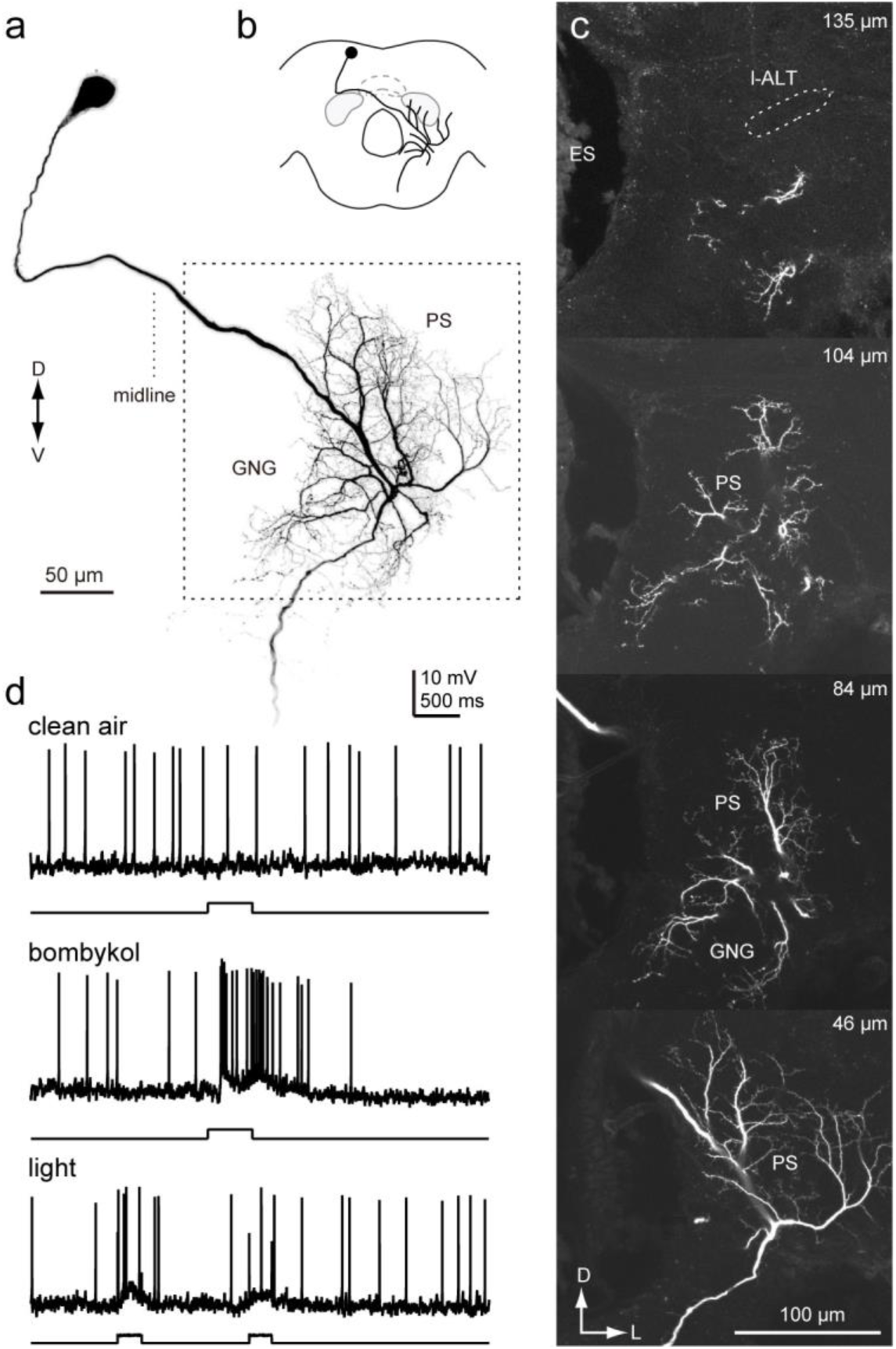
Morphology of a descending neuron innervating the PS. (**a**) Maximum intensity projection of the DN innervation in the brain. The neuron has smooth processes in the posterior slope (PS) and gnathal ganglion (GNG), and varicose processes in the GNG. (**b**) Schematic of neurite innervation for the neuron shown in (a). (**c**) Confocal stacks for the DNs shown in (a). The depth from the posterior brain surface is shown in *top-right*. The neuron lacks the innervation in the LAL. ES, esophagus; l-ALT, lateral antennal-lobe tract.

We also identified DNs with direct contact with the optic lobe (Figs. 11–13). All of these DNs innervated the inner lobula. We did not find DNs innervating the outer lobula or other parts of the optic lobe. Figure 11 demonstrates the morphology of a DN, which has smooth processes in the inner lobula and PLP, but not in the PS. The axon travels to the contralateral hemisphere via the LAL commissure and supplies varicose processes in the PS and GNG. We refer to this DN as lobula descending neuron 1 (LDN1). We were successful in staining a few more DNs with similar morphology to the neuron shown in Figure 11b. A recording was obtained from one example (#3); the DN did not show an obvious response to bombykol stimuli.

**FIGURE 11.**
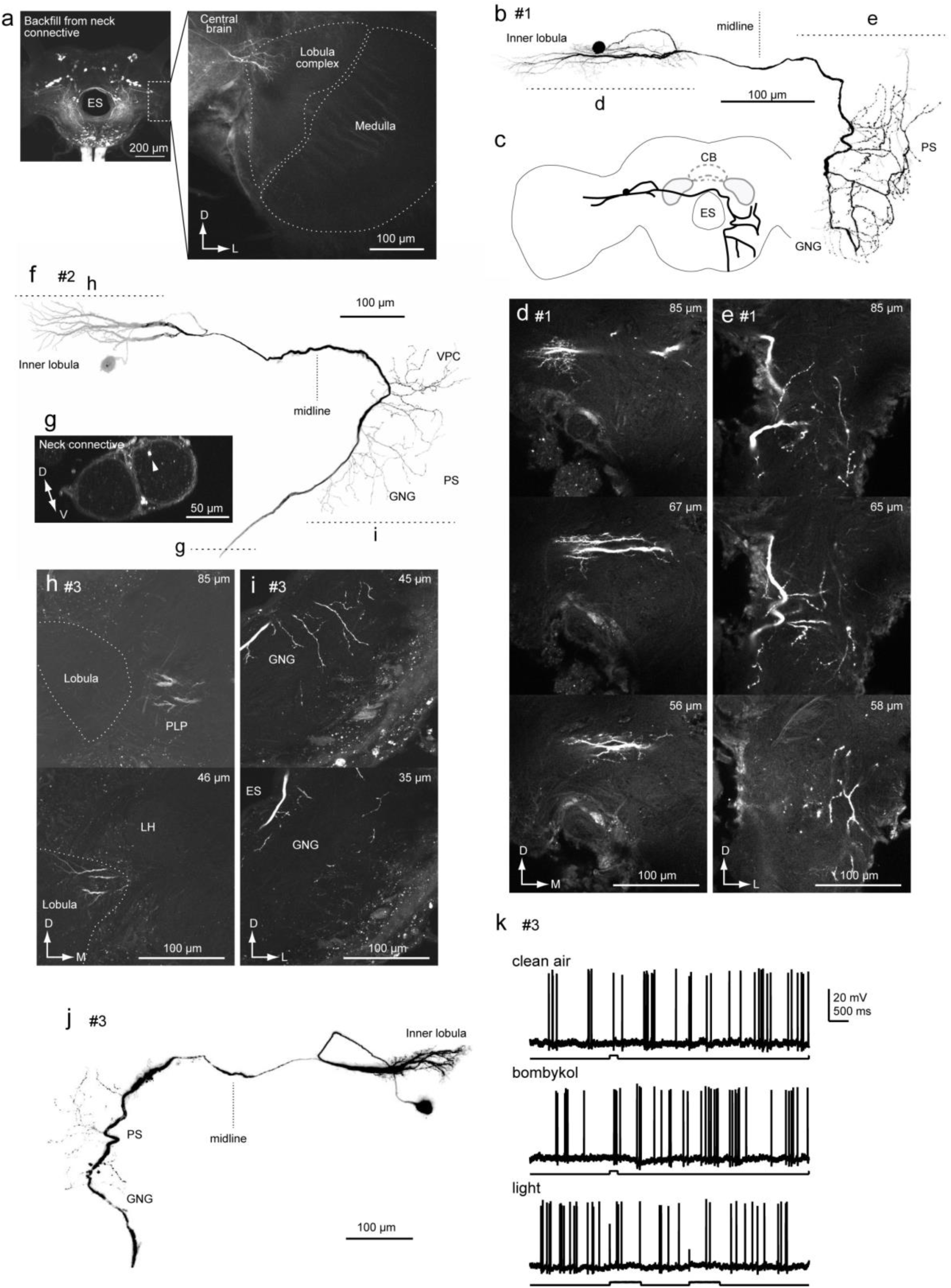
Lobula descending neuron 1. (**a**) An example of backfilling labels neurite entering into the optic lobe. Whole brain image (*left*) and high magnification of the optic lobe image are shown (*right*). (**b**) Morphology of a DN innervating the inner lobula. The DN has smooth processes in the inner lobula and posterior lateral protocerebrum (PLP) of the ipsilateral hemisphere and varicose processes in the posterior slope (PS) and gnathal ganglion (GNG) of the contralateral hemisphere. (**c**) Schematic of the neuronal morphology. (**d,e**) Confocal stacks of neuronal innervation in the lobula (d) and PS (e). The depth from posterior brain surface is shown in *top-right*. (**f**) Another example of LDN morphology. Basic anatomical feature is the same to the DN shown in (a). The position in the neck connective (**g**), innervation of smooth process (**h**) and varicose process are shown (**i**). (**j**) Third example of LDN morphology. (**k**) Response property of the neuron shown in (**j**). No obvious response was observed for the presentation of bombykol.

Figures 12 and 13 provide more examples of DNs innervating the inner lobula. The DNs have smooth processes in the inner lobula, LAL, PS, and SMP of the ipsilateral hemisphere, and varicose processes in the PS and GNG of the contralateral hemisphere. We refer to this DN type as lobula descending neuron 2 (LDN2).

**FIGURE 12.**
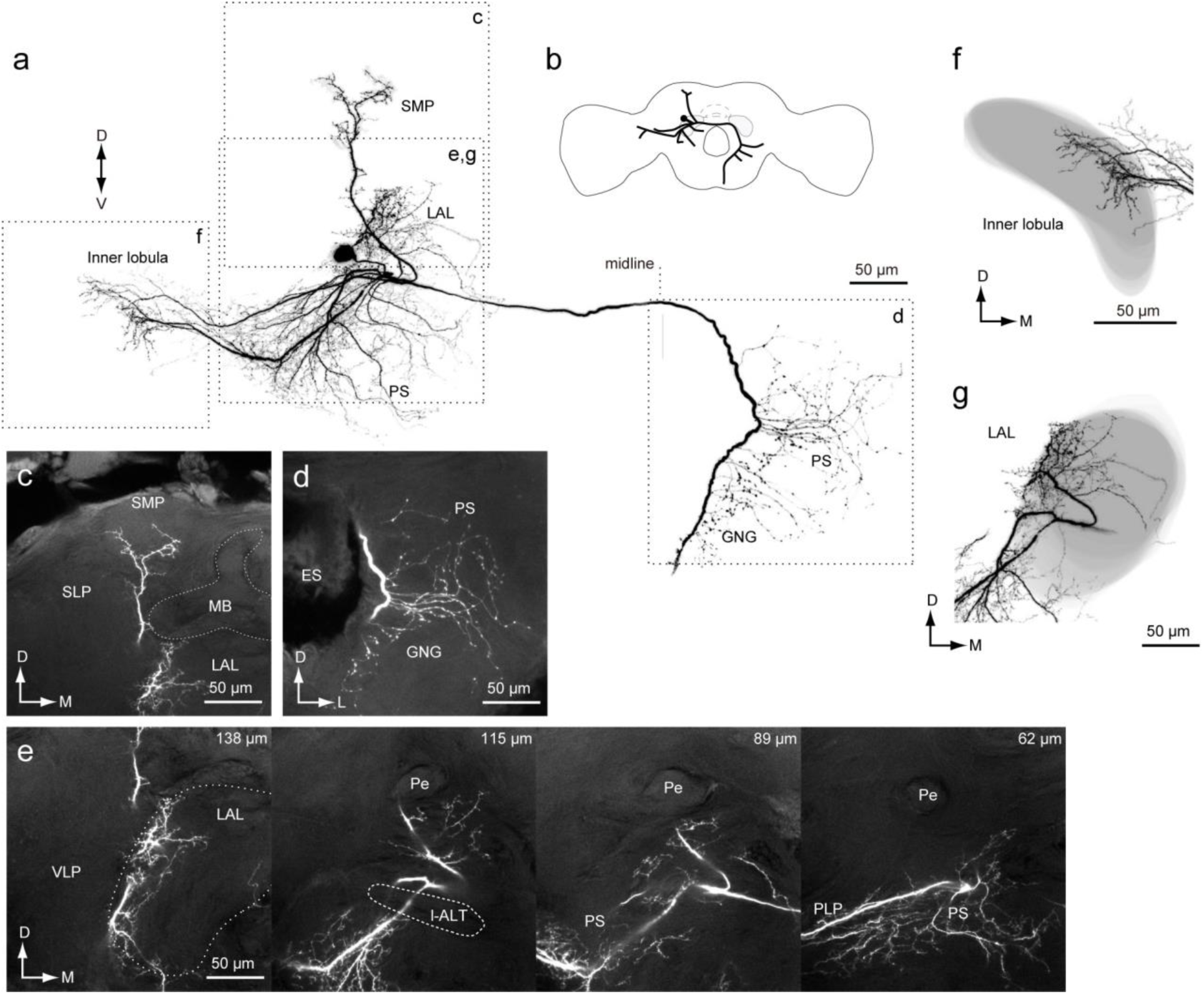
Lobula descending neuron 2: DN with wide field innervation in the brain. (**a**) Neuronal morphology in the brain. The neuron has smooth processes in the inner lobula, posterior lateral protocerebrum (PLP), posterior slope (PS), superior medial protocerebrum (SMP), ventral lateral protocerebrum (VPC) and the lateral accrssory lobe (LAL) of the ipsilateral hemisphere and varicose processes in the PS and gnathal ganglion (GNG) of the contralateral hemisphere. (**b**) Schematic of the neuronal morphology. (**c-e**) Confocal stacks for the innervation to the SMP (c), PS (d) and LAL (e). The depth from the posterior brain surface is shown in *top-right* (e). (**f,g**) Reconstruction of the neuropil shape and neuronal innervation in the inner lobula (f) and LAL (g). The innervation is biased toward the medial side in the inner lobula (f) and biased toward the lateral side in the LAL (g).

**FIGURE 13.**
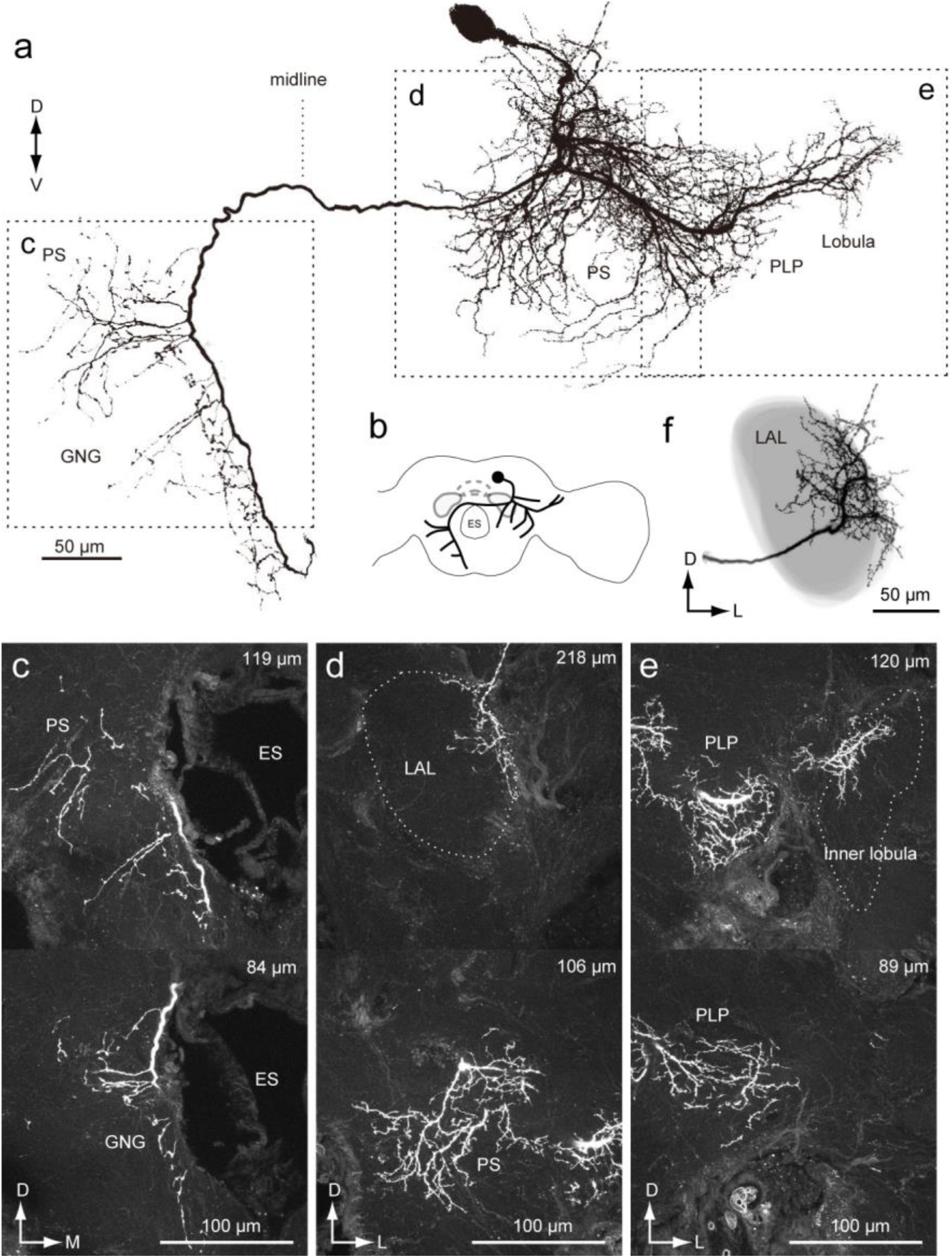
Lobula descending neuron 2: DN with wide field innervation in the brain. (**a**) Neuronal morphology in the brain. The neuron has smooth processes in the inner lobula, posterior lateral protocerebrum (PLP), superior medial protocerebrum, posterior slope (PS), ventral lateral protocerebrum (VPC) and the lateral accrssory lobe (LAL) of the ipsilateral hemisphere and varicose processes in the PS and gnathal ganglion (GNG) of the contralateral hemisphere. The morphology is similar to the neuron shown in Fig. 14. (**b**) Schematic of the neuronal morphology. (**c-e**) Confocal stacks for the innervation to the GNG (c), LAL and PS (d) and PLP and inner lobua (e). The depth from the posterior brain surface is shown in *top-right*. (**f**) Reconstruction of the LAL shape and neuronal innervation. The innervation is biased toward the lateral side of the LAL.

### 2.5. Biased innervation in the LAL

In studies prior to this definition, only four types, group-IA, IB, IIA and IID, were identified as innervating the LAL. In the present study, we have identified some novel DN types which innervate the LAL (group-IIE, IIF and LDN2; Figs. 7, 12 and 13). We analyzed the detailed morphology of neurite innervation by reconstructing LAL volume. The operational definition for the anatomical boundary of the LAL is defined by Iwano et al. ^22^. Using this definition, we re-examined their morphology. In the present study, we identified LAL innervation by group-IC and IB (Fig. 7c, e). We observed that the smooth processes are located above the depth of the saddle point of the l-ALT, which is classified as a part of the LAL under the definition ^22^. The newly identified group-IIF was seen to innervate the whole area within the LAL, similar to the innervation pattern for group-IID (Fig. 7d). We did not find the innervation in the LAL for group-IIC and - IIE. DNs innervating the LAL identified so far show innervation biased toward the medial side (lower division; Fig. 14a-h). Unlike the other types, LDN2 innervation is biased toward the lateral side of the LAL (upper division; Fig. 14i).

**FIGURE 14.**
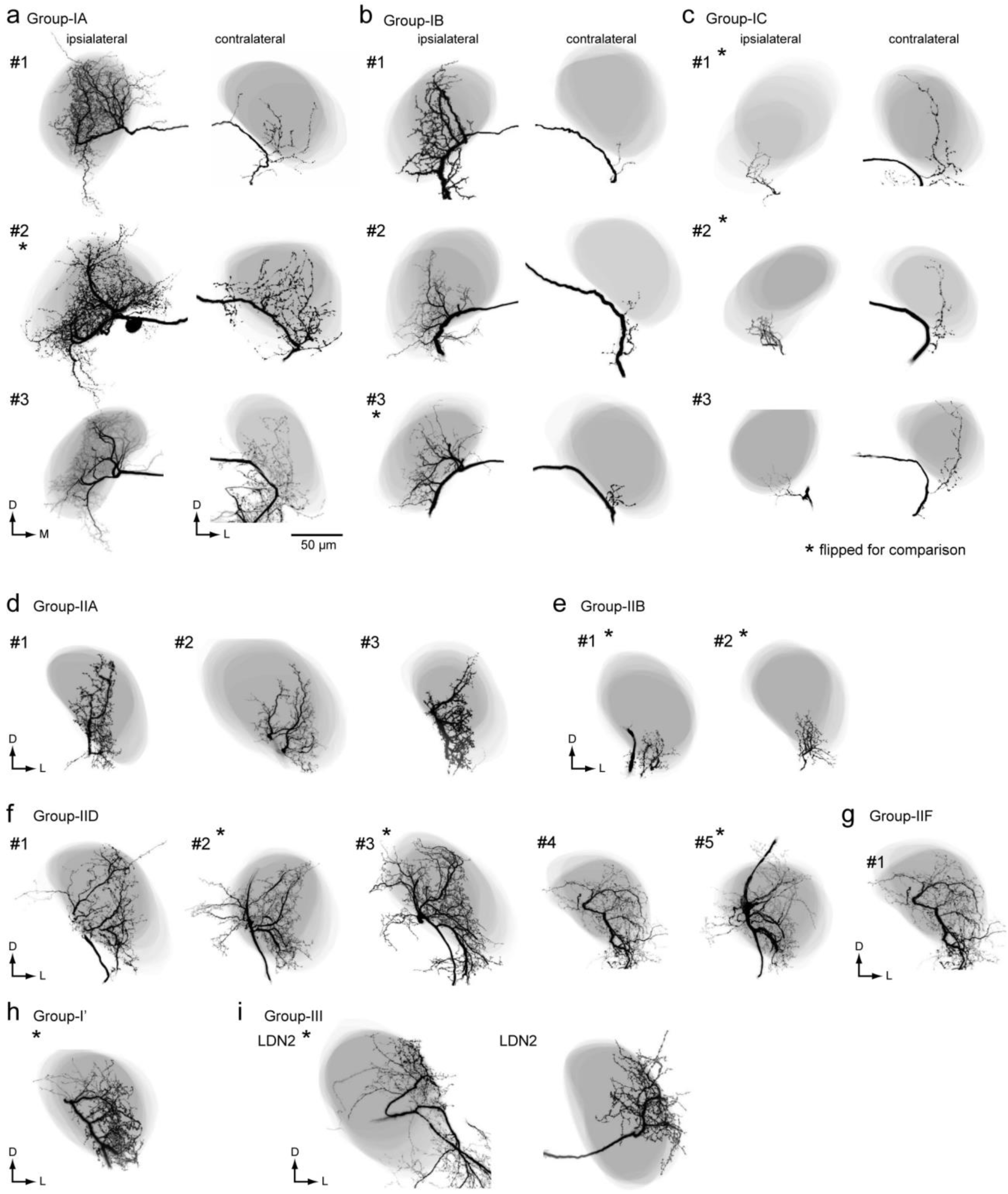
Innervation to the LAL by DNs. Neuronal innervation (black) and the LAL are shown (gray) for group-IA (**a**), group-IB (**b**), group-IC (**c**), group-IIA (**d**), group-IIB (**e**), group-IID (**f**), group-IIF DNs (**g**), group-I’ DN (**h**) and lobula DNs (LDN, #1 for neuron in Fig. 12, #2 for neuron in Fig. 13) (**i**). In some cases, flipped images are shown for the purpose of comparison (asterisk). Group-IIC and IIE DNs do not have innervation to the LAL.

## 3. Discussion

We found the DNs innervating the LAL also have innervation in the PS (Figs. 4, 7 and 8) and DNs arising from the PS which are responsive to the sex pheromone (Figs. 9 and 10). The functional difference of these two pathways, i.e., LAL+PS output vs. pure PS output would be interesting.

### 3.1. DN innervation to the LAL and PS

The strategy of moth pheromone orientation is composed of two behavioral modules: (1) a surge after the stimulus onset, and (2) casting after stimulus cessation ^23–27^. Flip-flop firing activity which is toggled by pheromone presentation, is thought to mediate command for zigzagging during the casting phase ^13^. The LAL is thought to be crucial generating flip-flop neural response in the silkmoth ^11,21^. A total of eight types of DNs (group-IA, IB, and IC; group-IIA, IIB, IID, and IIF; and group-III LDN2) have been identified to innervate the LAL. We revealed that these DNs have smooth processes in both LAL and PS and there are no DNs which receive input only from the LAL. In the present study, we also found group-III DNs from the PS, without innervation from the LAL, which exhibited excitatory response to the sex pheromone (Figs. 9 and 10; Supplementary Fig. 10). We did not observed flip-flop activity, suggesting that these DNs are involved in initial surge, rather than casting. These observation provide a possibility about the organization of descending pathways for pheromone orientation: (1) group-III DNs innervating the PS mediate initial surge, and (2) group-I/II DNs exhibiting flip-flop response meditate casting behavior (Supplementary Fig. 12). Flip-flop DNs show biphasic response property and the time period of early phase is well matched to the timing of initial surge, there is a possibility that these DNs also contribute the signal for initial surge.

The silkmoth is known to use visual information for pheromone orientation ^28,29^. The locomotion pattern is modified by the presence of the optic flow, suggesting the integration of pheromone and visual information. Optic flow modulates turning angular velocity during surge. DNs innervating the PS anatomically connected with lobula plate output ^30,31^. The lobula complex are known to process optical flow information ^32^. Such neuronal connection between the lobula plate and the PS is present in *B. mori* ^21^. Because group-III DNs arising from the PS show phasic response, there is a possibility that these DNs mediate the visual modulation of surge response. Contrary to the surge, optic flow modulate turn duration during casting ^28^ and optomotor response is absent in casting phase, suggesting different interactions between optic flow and pheromone processing pathways for surge and casting. We found group-I and group-II DNs have innervation to the PS. The dendrite of these DNs is the candidate site for the integration of pheromone-triggered premotor information and optic flow from the PS signal during casting.

### 3.2. Potential homology of neurons across species

Systematic data for the individual neuronal morphology of DNs is available in *Drosophila* ^9^, enabling comparisons of neuroanatomy with *Bombyx*. Comparisons at single neuron level reveal similar morphological feature between these genera. The characteristic morphological features of group-II DNs in *Bombyx* are: (1) cell bodies belonging to the cluster are located on the anterior surface beside the anterior optic tubercle, (2) they descend the ipsilateral side of the neck connective, and (3) they innervate the PS and some innervate the LAL (Fig. 7). This cell group is present in the closely related moth species, *Manduca sexta* ^33^. A neuroanatomical study reports that cell groups with these morphological feature, specifically, a group of ipsilaterally descending neurons, all of which has smooth process in the PS (DNa01~DNa10) ^9,34^, are present in *Drosophila*. Confocal image stacks of the morphology are available on the Janelia FlyLight database (http://www.janelia.org/split-gal4). As in *Bombyx*, most DNs in this group have varicose processes in the GNG, and some have additional branches into the LAL (DNa01, a02, a03, a06, and a08) (Supplementary Fig. 13a) ^9^. Cell groups with a similar morphological profile are present in other species, including the cricket *Gryllus bimaculatus* ^35^, the locust *Schistocerca gregaria* ^36^, and *Drosophila* larvae ^8^.

The present study characterized the morphology of group-I DNs as follows: (1) cell bodies belonging to the cell cluster are located on the anterior surface beside the antennal lobe, (2) they descend the contralateral neck connective, and (3) they have smooth processes in the PS (Fig. 4). DNs with these morphological features are present in *Drosophila* (DNb01 and DNb02) (Supplementary Fig. 13b). For example, DNb01 shows a morphological similarity to Group-IA DN in *Bombyx*. The innervation, both in the ipsilateral and contralateral LAL, is biased toward the medial side (lower division) and the DN has varicose processes in the medial PS of the contralateral hemisphere. In contrast, DNb02 has smooth innervation of both medial and lateral side of the PS, and the area of innervation in the LAL is small, reminiscent of group-IC in *Bombyx*. Potential homologous neurons to group-I DNs are present in other insects, including VG3 in the locust, *Schistocerca gregaria* ^36^, and B-DC1(5) in the cricket, *Gryllus bimaculatus* ^37^. Although the innervation of the PS has not yet been analyzed in detail, it is innervated by the posterior protocerebrum, in addition to the LAL. These observations suggest the potential homology of DN morphology for group-I and group-II DNs among insect species. There is a possibility is that group-I and group-II DNs are components of the ground pattern of the insect nervous system ^38^.

We have identified a number of DNs that innervate the optic lobe (Figs. 11–13). DNs innervating the optic lobe are also present in flies (lobula descending neuron, DNp11) ^5,9,39^. In contrast to LDN1 and 2, which innervate the inner lobula, DNp11 has smooth processes in the basal layer of the outer lobula. Unlike LDN in *Bombyx*, DNp11 innervates the antenno-mechanosensory motor center. *Drosophila* may only have a single type of DN that specifically innervates the lobula ^5,9^; however, *Bombyx* has several, with at least two types of DN innervating the lobula. Anatomical studies using backfill staining failed to find innervation to the lobula in other hemimetabolous insects such as the cockroach and cricket ^6,7^. There is a possibility that the LDN is an evolutionary new system in the moth and fly.

We have described the anatomical organization of DNs in the brain and identified the relationship between the LAL and PS in *Bombyx*. Based on comparative neuroanatomy between the moth and fly ^9^, we propose some hypotheses concerning a common design for the anatomical organization of the descending pathway in the insect nervous system: (1) the PS is densely innervated by DNs, (2) most DNs supply output to the GNG, (3) LAL and PS innervation are associated, and (4) innervation within the LAL is biased toward the medial side. These are testable by anatomical methods and should be examined in other insects. The neuroanatomy discussed here contributes to the investigation of the neural mechanisms underlying moth olfactory behavior.

## 4. Materials and Methods

### 4.1. Experimental animals

*B. mori* were reared on an artificial diet (SilkMate 2S and PS; Nosan Corporation Life-Tech Department, Yokohama, Japan). Adult male moths were used 2–7 days after eclosion.

### 4.2. Olfactory stimulation

Synthetic (E,Z)-10,12-hexadecadien-1-o1 (bombykol), the principal pheromone component of *B. mori*, with a purity of >99%, as confirmed by gas chromatography, was dissolved in high-performance liquid chromatography-grade n-hexane. The odorant (5 μl solution) was applied to a piece of filter paper (1 × 2 cm) and inserted into a glass stimulant cartridge with a 5.5-mm-tip diameter. The distance between the filter paper and the cartridge exit was approximately 7 cm. We applied 10 ng of bombykol to the filter paper, which corresponds to the amount that induces transient bursting activity in projection neurons in the AL and reliably triggers behavioral responses. Air or the odor stimulus was applied to either side of the antenna, and the exit of the cartridge was positioned 1.5 cm from the antennae. Compressed pure air was passed through a charcoal filter into the stimulant cartridge, and each stimulus was applied at a velocity of 500 mL min−1 (approximately 35 cm s−1), nearly the same as the flow speed as that produced when moths flap their wings. The moths were exposed to the odor for 200 or 500 ms, after which an exhaust tube was placed on the opposite side of the stimulant cartridge and the odor was removed (inner diameter, 4.5 and 15 cm from the antennae; ~55 cm s−1). All stimulant cartridges were sealed with a Teflon sheet, stored at −20 °C, and brought to room temperature prior to the recording session.

### 4.3. Intracellular labeling of single neurons

The staining procedure was conducted as previously described ^40^. After cooling (4°C, ∼30 min) to induce anesthesia, the abdomen, legs, wings, and dorsal side of the thorax were removed. The moth was fixed in a plastic chamber, and its head was immobilized using a notched plastic yoke slipped between the head and thorax. The brain was exposed by opening the head capsule and removing the large tracheae, and the intracranial muscles were removed to eliminate brain movement. The AL was surgically desheathed for inserting a microelectrode.

Filamented glass capillaries (TW100F-3; World Precision Instruments, Sarasota, FL, USA) were pulled on a micropipette puller (P-97 or P-2000; Sutter Instruments, Novato, CA, USA) and filled with 5% Lucifer yellow CH (LY) solution (Sigma, St. Louis, MO, USA) in distilled water or 1 M lithium chloride for staining neurons. The resistance of the electrodes was ∼60–300 MΩ. The electrodes were inserted using a micromanipulator (Leica Microsystems, Wetzlar, Germany), and a silver ground electrode was placed on the head cuticle. The brain was superfused with saline solution containing 140 mM NaCl, 5 mM KCl, 7 mM CaCl2, 1 mM MgCl2, 4 mM NaHCO3, 5 mM trehalose, 5 mM N-tris(hydroxymethyl)methyl-2-aminoethanesulfonic acid (TES), and 100 mM sucrose (pH 7.3). The incoming signals were amplified (MEZ-8300; Nihon Kohden, Tokyo, Japan), monitored with an oscilloscope (VC-10; Nihon Kohden), and recorded on a DAT recorder (RD-125T; TEAC, Tokyo, Japan) at 24 kHz. The acquired signals were stored in a computer using an A/D converter (PCI-6025E; National Instruments, Austin, TX, USA).

We stained each neuron using an iontophoretic injection of LY with a constant hyperpolarizing current (approximately −1 to −5 nA) for 1–3 min. After staining, the brain was superfused with saline solution containing 200 mM sucrose. Brains were fixed in 4% paraformaldehyde for 1–24 hours at 4°C. Brains were then dehydrated with 70%, 80%, 90%, 95%, and 100% ethanol (10 min in each) and cleaned in methylsalicylate for at least 30 min.

### 4.4. Backfill labelling

The ventral part of the neck was dissected to expose the neck connective. The nerve was stained by filling with saturated LY dissolved in distilled water from the cut end of the neck connective overnight at 4°C. The neck connective was placed in a pool made with Vaseline on a cover glass. To only stain one side, one of the neck connectives was damaged using forceps. After backfilling, the head was immediately removed. The brain was dissected from the head, and the brains were dissected, fixed, dehydrated, and cleared, as described above.

### 4.5. Imaging

Each stained neuron was visualized using a confocal imaging system (LSM510; Carl Zeiss, Jena, Germany) with ×40 (numerical aperture = 1.0) objective. LY-stained neurons were examined at a 458-nm excitation wavelength with a long-pass emission filter (>475 nm) in whole mounts. In some cases (Supplementary Fig. 6), we detected an autofluorescence signal using a HeNe laser at 543 nm and measured it with a 560-nm long-pass filter. Serial optical sections were acquired at 0.7- or 1.4-μm intervals throughout the depth of the neuron, and three-dimensional reconstructions of the labeled neurons were created from these sections.

### 4.6. Anatomical nomenclature

Regarding the neuroanatomical terminology, we followed the brain nomenclature proposed by Ito et al. ^17^. We updated the use of terminology in *B. mori*: we used the term “esophagus” for “oesophagus,” “gnathal ganglion” for “suboesophageal ganglion,” and “antennal-lobe tract” for “antenno-protocerebral tract.”

We had previously identified three types of DNs in the silkmoth brain ^11^ (Fig. 1). There are two groups of cell bodies on the anterior brain surface (Fig. 1a). Dorsal and ventral groups are termed “group-I” and “group-II”, respectively. Other types whose cell bodies were located on the posterior surface were classified as “group-III” (Fig. 1b). Group-I and group-II DNs descend via the contralateral and ipsilateral neck connective, respectively (Fig. 1d). Group-III contains both ipsilateral and contralateral DNs.

The anatomical border on the anterior side of the LAL is relatively well-defined and the posterior side was defined by the depth of the saddle point of the lateral antennal-lobe tract in most cases ^22^. Adjacent to the LAL is a small region called the VPC. We have not yet identified any local interneurons confined to this region and some LAL interneurons have additional branches within the VPC ^22^. The PS is unstructured neuropil, which is located posterior to the VPC. No clear anatomical boundary is available, except for the posterior border, which is defined by the brain surface. These three neuropils, LAL, VPC, and PS, are arranged serially and do not have clear anatomical boundary (Fig. 1e).

### 4.7. Data analysis

The segmentation and volume rendering of neurons and neuropils were performed in AMIRA 6.2 (FEI, Hillsboro, OL, USA). For images of single neuron morphology, masked images were used for visualization. We performed segmentation of individual neurons in confocal stacks. We first detected the signal with the Amira “Interactive Thresholding” function. We subsequently corrected any false detection by manual tracing. Using this image as a mask, we obtained the final masked images shown in the figures using a custom-made program written in MATLAB and the image processing toolbox (MathWorks, Natick, MA, USA). Maximum intensity projection images were prepared with ImageJ (National Institutes of Health, Bethesda, MD, USA) ^41^. The contrast and brightness of images were modified in Image J. Figures were prepared in Adobe Illustrator CS (Adobe Systems, San Jose, CA, USA).

## Acknowledgments

We thank Chika Iwatsuki, Satoshi Wada, Evan S. Hill, and Yoichi Seki for technical assistance; Shigeru Matsuyama for the purification of pheromone reagent. We are grateful to the FlyCircuit database from the NCHC (National Center for High-performance Computing and NTHU (National Tsing Hua University. This study was supported by Grant-in-Aid for Scientific Research from the Ministry of Education, Culture, Sports, Science, and Technology of Japan (Grant number: 17H05011 and 16H06732 to SN and 15H04399 to R.K.), Post-K Research and Development (R&D) projects (hp170227) to R.K. and Narishige Zoological Science Award to S.N.

